# Brain oscillations and connectivity in autism spectrum disorders (ASD): new approaches to methodology, measurement and modelling

**DOI:** 10.1101/077263

**Authors:** K. Kessler, R. A. Seymour, G. Rippon

**Affiliations:** Aston Brain Centre, School of Life and Health Sciences, Aston University, Birmingham, B4 7ET.

**Keywords:** Autism, Brain Oscillations, Gamma, Brain Connectivity, Predictive Coding

## Abstract

Although atypical social behaviour remains a key characterisation of ASD, the presence of sensory and perceptual abnormalities has been given a more central role in recent classification changes. An understanding of the origins of such aberrations could thus prove a fruitful focus for ASD research. Early neurocognitive models of ASD suggested that the study of high frequency activity in the brain as a measure of cortical connectivity might provide the key to understanding the neural correlates of sensory and perceptual deviations in ASD. As our review shows, the findings from subsequent research have been inconsistent, with a lack of agreement about the nature of any high frequency disturbances in ASD brains. Based on the application of new techniques using more sophisticated measures of brain synchronisation, direction of information flow, and invoking the coupling between high and low frequency bands, we propose a framework which could reconcile apparently conflicting findings in this area and would be consistent both with emerging neurocognitive models of autism and with the heterogeneity of the condition.

**Highlights:**

- Sensory and perceptual aberrations are becoming a core feature of the ASD symptom prolife.
- Brain oscillations and functional connectivity are consistently affected in ASD.
- Relationships (coupling) between high and low frequencies are also deficient.
- Novel framework proposes the ASD brain is marked by local dysregulation and reduced top-down connectivity
- The ASD brain’s ability to predict stimuli and events in the environment may be affected
- This may underlie perceptual sensitives and cascade into social processing deficits in ASD

## Introduction

Following recent changes in the classification of mental disorders, autism and autism-like disorders have been subsumed into a single spectrum of behaviours, Autism Spectrum Disorder (ASD). Although atypical social behaviour remains a key characterisation of ASD, the presence of sensory and perceptual abnormalities has been given a more central role. Indeed, (Baum, Stevenson, & Wallace, 2015) in a recent review suggested: “sensory processing is not only an additional piece of the puzzle but rather a critical cornerstone for characterising and understanding ASD”. It has additionally been suggested that the cascading consequences of low-level sensory and perceptual dysfunction could additionally present as various forms of social impairment (Freeman & Johnson, 2016; Lawson et al., 2014; Schilbach, 2016). An understanding of the origins of these low-level atypicalities could thus prove a fruitful focus for ASD research.

A decade ago, emergent theories of the role of gamma band activity (GBA) in ‘temporal binding’, in other words the formation of a coherent perceptual result (“percept”) essential for accurate information processing, indicated that gamma could be a useful ‘candidate’ frequency for characterising the cortical correlates of sensory and perceptual atypicalities in ASD (Rubenstein & Merzenich, 2003). Abnormal GBA in ASD was linked to models of excitation-inhibition imbalance and atypical cortical connectivity (Rippon et al., 2007). Subsequent research focussed on measures of GBA in ASD, particularly in association with visual and auditory processing but also, more recently, considering resting-state data. An overview of this research shows a lack of consistency, with no clear-cut picture emerging of the nature of the local dysregulation that would be predicted from the sensory and perceptual difficulties evident in ASD. We will demonstrate that this may partly be due to the use of different measures of GBA, but also to the inadequacy of early gamma metrics or an overly simplistic focus on a single frequency measure. Research into cortical connectivity showed a similar lack of consensus, although there have been claims as to ‘firm findings’ of long-distance hypoconnectivity (Hughes, 2007). Again, as we will show, this is associated with the use of different connectivity metrics, as well as there being a focus on fMRI, inappropriate for investigation of more sophisticated temporal and spectral models of cortical connectivity that are now emerging. As noted by Vissers, Cohen, & Geurts, (2012) this is a field in need of refined models, methodological convergence and stronger behavioural links.

Recent research developments into gamma brain oscillations have marked increasing levels of sophistication in their measurement, modelling and interpretation, including greater complexity in analytical techniques, beyond a focus on within-band evoked or induced power responses. Measures of phase-synchrony and low-high cross-frequency coupling (CFC), together with quantification of gamma-based brain network properties, offer much more nuanced characterisation of both task-related and resting-state activation patterns. These can inform complex models of sensory and perceptual processing, such as Bayesian predictive coding (Pellicano & Burr, 2012) and feedforward/feedback propagation (Bastos et al., 2015; Samson et al., 2012) and thence allow testable predictions associated with the sensory and perceptual atypicalities associated with ASD.

The aim of this review is to track the progress to date in this field and identify possible sources of reconciliation. We propose a framework whereby anomalous GBA and deficient CFC processes will provide evidence of a local dysregulation of optimal processing associated with an excitation-inhibition imbalance. This will present as increased or decreased GBA depending on context and task. There will be disrupted signal-to-noise ratios due to the suboptimal balance between excitation and inhibition and consequent abnormalities in gamma feedforward connectivity. This feedforward dysfunction will disrupt the formation of long-range connectivity, reducing reciprocal feedback and top-down control, as measured by atypical global phase coupling across relevant brain areas. As a consequence, there will be an overall failure in optimal predictive coding of the environment (Arnal & Giraud, 2012), with associated atypicalities in sensation and perception (Friston, Lawson, & Frith, 2013; Pellicano & Burr, 2012). The metrics generated by this framework could be linked to the development of abnormal ‘protective’ behaviours (Van de Cruys et al., 2014) and consolidate the association between sensory and social symptomatology in ASD. Before discussing these new metrics and the proposed framework, we provide an overview of ASD symptomatology in the next Section.

## 1. Autism Spectrum Disorders

Autism Spectrum Disorder (ASD) is a highly heritable neurodevelopmental condition, with a prevalence of around 1 in 88 (Baio, 2012). The condition is characterised by persistent deficits in social communication and interaction as well as restricted and repetitive patterns of behaviour, interests, or activities (APA, 2013). It is also a highly heterogeneous condition (Jeste & Geschwind, 2014) with widely varying levels of intellectual ability and degrees of symptom severity, as well as low levels of co-occurrence of what are claimed as the core impairments (Happé & Frith, 2006). This behavioural heterogeneity is a key challenge to any attempt to identify the underlying causes of the condition.

Although atypical social behaviour remains a primary characterisation of ASD, the presence of sensory abnormalities has recently been given a more central role, consistent with reports that over 90% of ASD individuals experience some form of sensory abnormality in proprioception, olfaction, auditory and/or visual domains (Hazen et al., 2014; Leekam et al., 2007). Such problems have been described in qualitative interviews (Kirby, Dickie, & Baranek, 2015), as well as using self-report questionnaires such as the Sensory Over-Responsivity Scale (Baranek et al., 2006; Liss et al., 2006) and the Glasgow Sensory Questionnaire (Robertson & Simmons, 2013). ASD individuals commonly report a severe hyper-sensitivity to arousing stimuli, although hypo-sensitivity is also reported for a subset of individuals (Ben-Sasson et al., 2009). A recent review has drawn attention to the potential that increased insights into ASD sensory problems could bring; not only to understanding (and possibly alleviating) the ASD experience, but also to measuring and mapping the neural underpinnings of ASD (Baum et al., 2015).

Unusual aspects of perceptual processing are also evident in ASD. These are commonly characterised as the tendency of ASD individuals to focus on local detail at the expense of global processing (Bölte et al., 2007). Indeed, Kanner’s original profiling of autism noted that his patients often “failed to experience wholes despite paying attention to the constituent parts” (Kanner, 1943). This is the converse of typical perceptual processing, where stages are temporally organised so that they proceed from global structuring towards increasingly fine-grained analysis. This local bias in ASD has been shown to manifest itself via sharper discrimination thresholds in response to luminance gratings, auditory tones and tactile cues (O’Riordan & Passetti, 2006; Simmons et al., 2009) as well as enhanced performance on visual search (O’Riordan, 2004) and embedded figures tasks (Happé, 1999; though see White & Saldaña, 2011). This focus on local detail in ASD is often accompanied by apparent deficits in global processing. For example, participants with ASD are less susceptible to the perception of illusory figures like Kanisza triangles, which requires the automatic integration of contextual information (Bölte et al., 2007; Walter, Dassonville, & Bochsler, 2009) into a so-called global “gestalt”, and are slower on the hierarchical figures task, which requires a mapping between local and global levels of processing (Scherf et al., 2008). A local bias will also disrupt task performance where integration of local parts into a global whole or gestalt is required (Bölte et al., 2007; Dakin & Frith, 2005). Interestingly a recent meta-analysis of perceptual processing in ASD suggests that global processing may be temporally slower in ASD, meaning that ASD individuals rely on local processing strategies to a much greater extent than typically developing controls (Van der Hallen et al., 2015).

It has been suggested that sensory and associated perceptual difficulties may also underpin the patterns of restricted interests and activities typical of ASD and could even cascade into the characteristic social and behavioural deficits during development (Behrmann et al., 2015). Anomalous individual sensory and perceptual experiences could well render the world “painfully intense” leading to social withdrawal and/or obsessive focus on deliberately limited experiences (Markram & Markram, 2010) or cause difficulties with the downstream integration of the complex information necessary for responding appropriately to social rules (Gepner & Féron, 2009). An understanding of the mechanisms underlying these sensory and associated perceptual atypicalities could thus prove a fruitful focus for ASD research. The novel framework that will be proposed in Section 6 aims at explaining sensory aberrations and their knock-on effects on higher-level cognitive processing in ASD. It is based on deficits in feedforward-feedback brain mechanisms as reflected by anomalies in brain oscillations and their interplay across frequency bands and between brain areas. We will therefore introduce a few basic concepts in the following sections that will facilitate understanding of the remainder of the manuscript.

## 2. Brain oscillations, sensory-perceptual processing and cortical connectivity

Successful processing of sensory information by the brain requires a mechanism that can effect the integration of separate parts into coherent wholes, via the synchronisation of specialised neural networks in the brain. A ‘temporal binding’ mechanism has been proposed that creates and maintains the transient neuronal assemblies underpinning the integration of information necessary for perception (Singer & Gray, 1995), and could also serve as a general mechanism of inter-cortical information transmission both locally and distally (Varela et al., 2001). There is strong evidence that cortical oscillations are involved in this process (Fries, 2005), in particular oscillations in the gamma-band (<40Hz) (König, Engel, & Singer, 1995; Singer & Gray, 1995).

### 2.1 Feature integration and gamma band activity (GBA)

Research has suggested that gamma-band *synchrony* is a plausible mechanism to bind groups of spatially separable neurons, each encoding specific aspects of a stimulus, into a coherent whole (Singer & Gray, 1995). In this so-called Binding by Synchrony (BBS) account, gamma synchrony is hypothesised to determine optimal neuronal response timing (Buzsáki & Wang, 2012; Whittington et al., 2011) and ensure maximum accuracy in stimulus processing. A distinct, but related hypothesis is that gamma-band synchrony allows the efficient exchange of information between neurons at both the local and global scales (Fries, 2005). By oscillating at high frequencies, a neuron’s window of excitability becomes constrained to distinct periods of the gamma cycle (Tiesinga et al., 2004). Only neurons sending and receiving input in a temporally synchronised manner, such that periods of pre and post-synaptic excitability align, are thought to be able to communicate. This is hypothesised to render neural communication precise and effective (Bastos, Vezoli, & Fries, 2015); not only during sensory processing but across multiple cognitive domains.

Given the hypothesised functional role of gamma in the controlling the timing of cortical responsivity, both locally and distally, there has been a focus on ways of quantifying temporal synchronicity in GBA. For example, based on a method developed by (Lachaux et al., 1999), it is possible to measure synchronisation between different cortical areas of GBA phase, relatively independent of amplitude. This is known as the Phase-Locking Factor or Value (PLF or PLV), with values between 0 and 1 (with 1 as maximum synchrony), and gives a measure of the percentage of measured signals in phase across trials or periods of measurement. PLF can also be applied to measures of phase synchrony between pairs of signals for a given frequency (e.g. Martini et al., 2012). Phase consistency across trials (not across brain areas) can also be measured via phase coherence indices that have been variously termed as Inter-Trial Coherence (Port et al., 2007), Inter-Trial Phase Coherence and/or Phase Synchrony (Isler et al., 2010). While these measures are a significant improvement over mere power measures, giving insight into presence or absence of systematic rhythms across repetitions (trials), veridical phase estimates in gamma frequency, especially in high gamma, are hard to achieve due to the large spread of the respective frequency bands (e.g. 30-60, 60-90). This might result in fluctuations across studies in terms of which frequencies reveal significant phase alignment effects. This further applies to gamma phase-relationships between brain areas. In contrast, recently proposed measures of local and global systematicity of processing such as cross-frequency coupling (CFC), especially in the form of phase-amplitude coupling (PAC), might not suffer from erroneous gamma-phase estimates and will be discussed in the next section.

### 2.2 Stimulus representation, predictive coding and cross-frequency coupling (CFC)

GBA has been described as the basis of a ‘temporal code’ which can, for example, exactly specify stimulus features for memory matching (Herrmann, Fründ, & Lenz, 2010), with synchronisation or desynchronisation serving to ‘sharpen’ or more closely specify stimulus representation (Moldakarimov, Bazhenov, & Sejnowski, 2010). GBA elicited by sensory input will feed forward to higher brain areas to inform higher-level encoding and processing (Bastos et al., 2015).

Contemporary models of perception posit a Bayesian model of predictive coding where top-down hypotheses, prior expectations or predictions are matched to bottom-up input from sensory areas (Friston, 2005). Discrepancies are known as prediction errors and will result in alterations to current predictions. Prediction errors can be minimised by accurate matching of input to expectation but can be maximised by deficient or over-specific predictions and/or inaccurate bottom-up sensory coding, for example, a poor signal-to-noise ratio (SNR) in brain activity at the sensory level (Moldakarimov et al., 2010). An optimal balance in SNR including top-down regulation in form of selection and filtering allows for optimal predictive coding of the environment. With intact top-down regulation (filtering, selection, integration) local encoding can be predictive for “what” should happen and “when” (Arnal & Giraud, 2012), thus mainly processing deviations from predictions, resulting in a system that is highly efficient and proactive.

Electrophysiologically, predictive coding approaches have been linked to cross-frequency coupling (CFC) of specific brain oscillations (Arnal & Giraud, 2012), with high-frequency gamma oscillations proposed to play a prominent role in the coding of the prediction error, i.e. the signal that reflects the deviation between sensory input and prediction, and lower frequencies related to top-down establishment of predictions. The use of phase amplitude coupling (PAC) metrics has provided further insights into these partnerships (see Figure 1). Phase-amplitude coupling is the mechanism where the phase of a lower frequency oscillation in one area (theta, alpha, beta) has been shown to modulate the amplitude of higher frequency oscillations (commonly gamma) in the same or other areas (Canolty & Knight, 2010). The efficiency of this coupling is taken as a measure of the functional connectivity between the various sources; both long- and short-range connectivity can be studied using this approach (Hyafil et al., 2015; Palva & Palva, 2011; Varela et al., 2001; but see modelling results by Peterson & Voytek, 2015) for a possible caveat). As PAC can measure the efficiency of interactions within neuronal populations operating at high and low frequencies, it can also be taken as an optimal measure for local processing efficiency. Recently, computational modelling of oscillatory activity indeed suggested that PAC may be a key component in balancing excitation-inhibition interactions and maximising information flow between brain areas (Onslow, Jones, & Bogacz, 2014; Peterson & Voytek, 2015).

**Figure 1.**
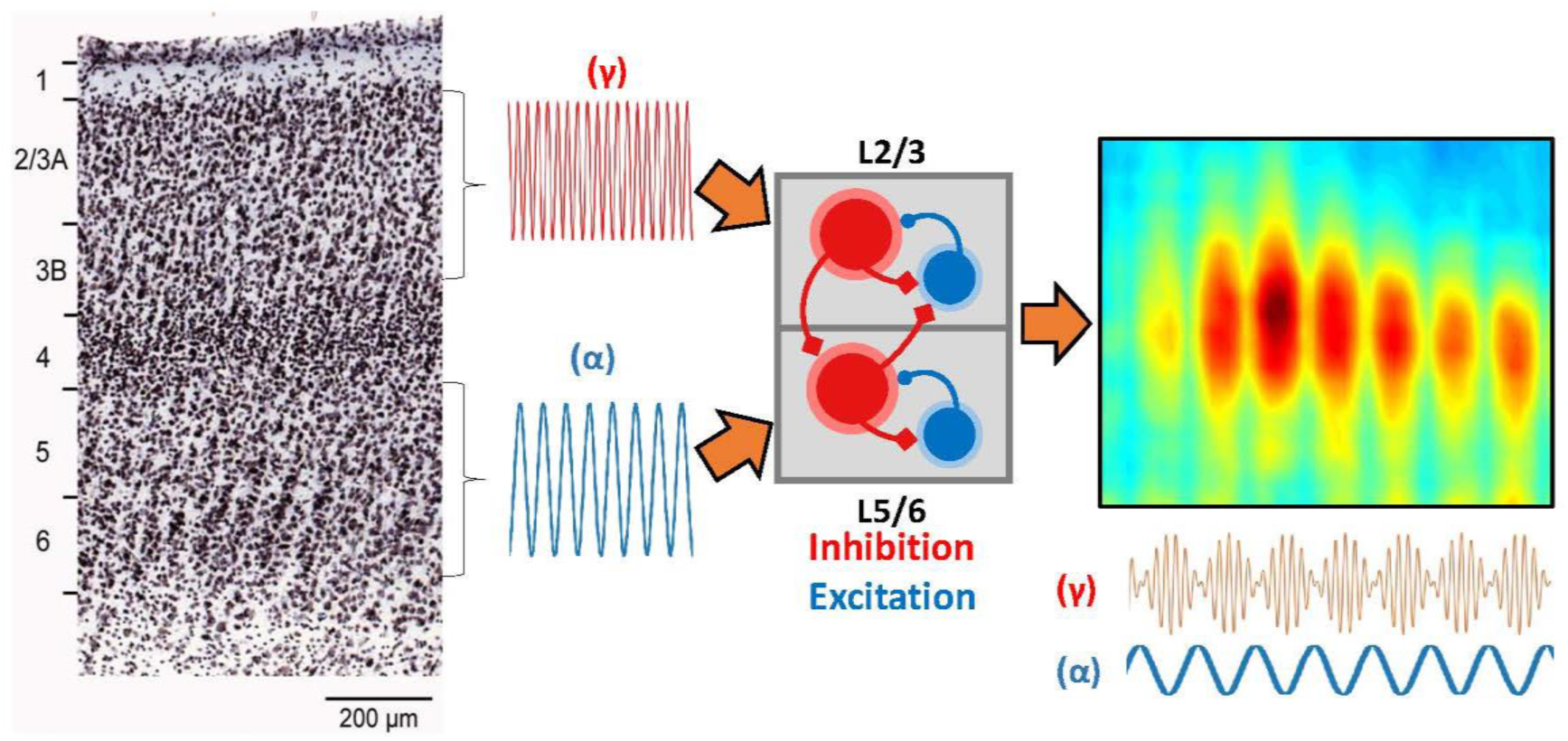
Illustration of the emergence of alpha (α) – gamma (γ) phase-amplitude-coupling (PAC) from the cerebral cortex. Gamma (30-80Hz) and alpha (8-13Hz) oscillations have been shown to emerge separately from supragranular layers (2/3) and superficial layers (5/6) of cortex. Meijas et al. (2016) propose that these rhythms interact via an inter-laminar coupling circuit based on interactions between excitation (blue) and inhibition (red). This results in the amplitude/power of the supragranular gamma rhythm becoming entrained to the phase of the superficial alpha. The photomicrograph of cortical layers has been reproduced from Markov et al., (2014).

PAC has been shown to be correlated with task difficulty and task performance (Canolty et al., 2006), with theta-gamma coupling closely linked to changes in memory state and memory performance (Canolty et al., 2006; Lisman & Jensen, 2013). Visual processing seems to elicit reliable PAC between the alpha & gamma bands (Voytek et al., 2010), with the phase of alpha oscillations thought to temporally segment visual gamma via inhibition (Bonnefond & Jensen, 2015). In auditory/speech models it has been shown that auditory cortical responses to speech occur in the theta range, which then modulates gamma activity (Giraud & Poeppel, 2012; Schroeder et al., 2008).

### 2.3 GBA, PAC and cortical connectivity

As above, early models of the role of brain oscillations as indices of cortical connectivity focussed on GBA given its ‘temporal binding’ role (Buzsáki & Wang, 2012; Tallon-Baudry & Bertrand, 1999; Whittington et al., 2011). Thus, where abnormal connectivity was implicated in brain disorders, the research focus was on GBA. More recently, where CFC has been proposed as a more detailed framework for modelling cortical connectivity, PAC measures have been employed to illustrate deficiencies. While these measures are optimally suited to investigate integrity of local connectivity, it has been suggested that long-range connectivity between cortical and thalamic systems is mediated by low frequencies that in turn may drive local gamma in form of PAC (e.g. Onslow et al., 2014; Simon & Wallace, 2016). Low-frequency connectivity has repeatedly been associated with predominantly top-down connectivity that implements the described selection, filtering and integration mechanisms (Engel, Fries, & Singer, 2001; Jensen et al., 2015) as well as the top-down signaling of predictions in terms of *what* is to be expected and *when* (Arnal & Giraud, 2012).

As atypical cortical connectivity has been suggested as a core feature in Autism Spectrum Disorders (Gepner & Féron, 2009), a major emphasis in research in this area has been, initially, on abnormal measures of GBA and, more recently, on measures of CFC, phase-based measures of long-range connectivity, and predictive coding. These will be reviewed in Sections 5.

## 3. Neurophysiological Models of ASD

As identified in Section 1, ASD is highly heritable (Betancur, 2011) but also extremely behaviourally heterogeneous (Jeste & Geschwind, 2014). Identifying a potential endophenotype for ASD is therefore complex, requiring the identification of some form of genetically mediated and quantifiable biological characteristic which could be linked to the diverse behavioural symptoms (Moseley et al., 2015). Whilst no common biological features have been identified, recent work has shown that many genetic mutations associated with ASD disrupt the development of typical synaptic structure and function (Chen et al., 2015; De Rubeis et al., 2014). A focus of the search for the neural underpinnings of ASD has therefore been on the resulting consequences of this synaptic dysfunction, such as atypical cortical organisation and its consequences for brain development and function. Such research involves animal and translational models of ASD at the cellular level as well as neurocognitive models matching symptom profiles to hypothesised neural phenotypes, such as aberrant cortical connectivity.

### 3.1 Excitation-inhibition in autism

Animal models of autism have demonstrated that in utero exposure to valproic acid (VPA) or targeted knock out models will damage inhibitory neurons linked to gamma –inducing inhibitory process (Sohal et al., 2009). This has been shown to significantly increase local connectivity (Rinaldi, Silberberg, & Markram, 2008), and produce a wide-range of autism-like symptoms (Gogolla et al., 2009). It has been proposed that this damage will affect the appropriate excitatory/inhibitory function in developing neural circuits by its effect on cortical minicolumns (Casanova et al., 2002; Rubenstein & Merzenich, 2003). These are neuronal ‘microcircuits’ which are distributed throughout the cortex, and comprise narrow radial columns of 80-100 neurons surrounded by gamma amino butyric acid - (GABA-) containing interneurons, which act to segregate individual minicolumns, regulating their output and ensuring discrete channels of intra-cortical communication. Synchronisation of these local systems is indexed by cellular activity around 40 Hz, in the gamma frequency range (Whittington et al., 2011). Dysfunction in these local inhibitory neurons, reflected by abnormalities in GABA levels, could cause a generic disruption in the excitation-inhibition balance within the cortex, with reduced inhibition as indexed by GABA levels associated with increased excitation. This would manifest as increases in high frequency or ‘noisy’ activity in the brain, common in ASD (Berg & Plioplys, 2012; Spence & Schneider, 2009), and associated anomalies in GBA. In this model, then, mini-column deficits would be associated with hyper-activity at the local level. This would have implications for the feedforward role of gamma in sensory processing, with the emerging disorganisation between local circuits disrupting the synchronisation necessary to ensure appropriate correlated co-ordination of stimulus processing (see Section 2), resulting in atypical sensory responses associated with abnormal gamma activity. Further, abnormalities in local circuits could impede the formation of long-distance connectivity between various, functionally specialised parts of the cortex, resulting in significant hypo-or under-connectivity cross-cortically (Casanova et al., 2002; Courchesne & Pierce, 2005; Lewis & Elman, 2008; Rubenstein & Merzenich, 2003). The model is now being supported at the cytoarchitectural level (Casanova et al., 2013; Casanova & Trippe, 2009; Stoner et al., 2014) with additional support from in vivo measures of GABA in ASD populations (Gaetz et al., 2014). However, a recent review of neurophysiological and psychophysiological research into alterations in the balance of excitation-inhibition in ASD has concluded that, while the evidence for an imbalance is strong, it may arise not only from excitation being increased relative to inhibition (as above) but also from increased inhibitory processes resulting in imbalance relative to excitatory processes (Dickinson, Jones, & Milne, 2016). It is possible that either or both types of imbalance could be associated with different manifestations of atypical behaviour in ASD individuals and thence underpin the problematic heterogeneity of the disorder.

### 3.2 Aberrant cortical connectivity in ASD

As above, it is proposed that the disruption of the excitation – inhibition imbalance at the cellular level has consequences for patterns of connectivity in the autistic brain. As outlined in Section 2, the synchronisation of specialised neural networks is key to normal sensory, perceptual and cognitive function. Disruption in neural synchrony or failures in dynamic network communication have been hailed as a unifying explanation for a wide range of behavioural and neurocognitive disorders and have been widely reviewed (Menon, 2011; Uhlhaas & Singer, 2006; Voytek & Knight, 2015). A key aspect of the development of these models has been the major advances in techniques for studying connectivity (see Hutchison et al., 2013). Earlier research employed functional magnetic resonance imaging (fMRI) techniques, with the fine spatial resolution enabling detailed mapping of network nodes and pathways, thus capturing the structural characteristics of hypothesised networks. Estimates of functional and/or effective connectivity between voxels or Regions of Interest (ROIs) were obtained using correlation or causal modelling metrics (Friston, 2011). However, fMRI is less able to capture the proposed temporal dynamics of activated networks (Logothetis, 2008), and cannot measure the spectral characteristics, including the gamma-band response (GBA), which appears to be a key index of neural synchronisation (Fries, 2005, 2015). For this, the millisecond temporal resolution offered by EEG and MEG is required (Lopes da Silva, 2013).

The study of neuronal circuit dysfunction specifically in ASD has formed a major part in connectivity research and has, similarly, been the subject of a number of reviews (Belmonte et al., 2004; Geschwind & Levitt, 2007; Hahamy et al., 2015; Müller et al., 2011; Rippon et al., 2007; Vissers, Cohen, & Geurts, 2012; Wass, 2011; Picci, Gotts, & Scherf, 2016). The consensus from *early* findings of task-related activity was of accumulating evidence of long-distance structural and functional underconnectivity between specialised cortical regions underpinning a wide-range of perceptual and cognitive processes (Hughes, 2007; Just, Cherkassky, Keller, Kana, & Minshew, 2007; Just, Cherkassky, Keller, & Minshew, 2004; Kana, Keller, Minshew, & Just, 2007; Koshino et al., 2008). However, Thai, Longe, & Rippon, (2009) noted that there were methodological differences between those fMRI studies whose findings supported a ‘General Underconnectivity’ hypothesis and those that did not, with the former more focussed on task-related regional interconnections and the latter more likely to take a whole-brain, task-free approach, often identifying regions of atypically increased functional connectivity in ASD. There is also accumulating evidence of atypical connectivity between subcortical and cortical structures in ASD (Cerliani et al., 2015), in particular networks involving the thalamus (Cerliani et al., 2015), and amygdala (Kleinhans et al., 2016). Additionally, a recent fMRI study using a naturalistic movie-viewing paradigm observed aberrant connectivity between several cortical regions and a ventro-temporol-limbic subcortical network involving the amygdala, striatum, thalamus and parahippocampal structures (Glerean et al., 2016).

More recent fMRI studies using connectivity measures derived from resting state measures report both under- (Dinstein et al., 2011) and over-connectivity (Keown et al., 2013; Supekar et al., 2013) or normal patterns (Tyszka et al., 2014). However, a recent article by (Hahamy et al., 2015) suggested that there are marked individual differences (or idiosyncrasies) shown by whole-brain analyses of connectivity, with ASD participants showing substantially more deviation from group-averaged patterns of inter-hemispheric connectivity. They noted that closer attention to this source of variation may resolve some of the apparent contradictions. Developmental variations and trajectories may also affect measures of connectivity in ASD – for example Uddin, Supekar, & Menon (2013) suggest that individuals with ASD show general hyperconnectivity in childhood, but fail to display characteristic increases in connectivity during adolescence through to adulthood. As will be discussed below (see Section 5), inconsistent results across EEG/MEG connectivity studies may reflect such developmental factors in GBA (Tierney et al., 2012) and also differences in the patterns of connectivity as reflected by different brain oscillation frequencies (Kitzbichler et al., 2015; Von Stein, Chiang, & König, 2000).

Thus far, studies of cortical connectivity have mainly reported on long-distance or inter-regional connections, principally determined by the imaging techniques employed. Measures of local connectivity require the temporal resolution appropriate to the transient nature of local processing and additionally access to the variations in the spectral characteristics indexing the activity of local systems. With respect to patterns of local connectivity, studies using EEG and MEG have identified patterns of cortical oscillations which would be consistent with localised hyper-reactivity and excitation-inhibition imbalance (Cornew et al., 2012; Orekhova et al., 2007) (but equally, other studies reporting results consistent with reduced connectivity at the local as well as the long-distance level e.g. Khan et al., 2013).

These will be reviewed in more detail in Section 5. One key issue to be considered is the validity of the spectral measures of connectivity being used, as inferences based on power measures alone can be inconsistent with more complex measures of coherence/phase-locking (Port et al., 2015) or of cross-frequency coupling (Canolty & Knight, 2010).

Currently then, although there is consensus that disruptions in neural synchrony or failures in dynamic network communication can be a “unifying explanation”, in neurocognitive disorders, with respect to ASD, the picture is far from consistent. A recent review by Picci et al, 2016 pointed out that several studies have reported connectivity patterns varying with symptom severity, thus suggesting that the ‘heterogeneity’ issue is also evident in the cortical as well as the symptom profiling of ASD and should be taken into account in exploring the possibility that measures of cortical connectivity might serve as a useful endophenotype in ASD (Moseley et al., 2015; Simon & Wallace, 2016).

## 4. Neurocognitive models of ASD

Research into the neural underpinnings of ASD has also focussed on core deficits of the condition and linked these to inferred underlying cortical processes. This has included models of the diagnostic impairments such as those in social communication and interaction (Simon Baron-Cohen, 1997) and repetitive and executive function (Russell, 1997), but more recently has considered the neural bases of atypical sensory and perceptual function.

### 4.1 Weak Central Coherence/Enhanced Perceptual Processing

Research into atypical sensory and perceptual processing in autism has inspired the development of theoretical models such as the ‘weak central coherence’ (Happé & Frith, 2006) and ‘enhanced perceptual functioning’ accounts (Mottron et al., 2006). The weak central coherence theory suggests that ASD individuals are merely ‘biased’ towards attention to fine-grained local detail and are less distracted by the context of the whole stimulus array. Frith has demonstrated that given the appropriate instruction, ASD individuals can perform normally on tests of global processing (Dakin & Frith, 2005), suggesting that local processing is more of a ‘default option’ for ASD individuals. Mottron et al., (2006) however, suggests that the evidence of superior performance in tasks such as visual search or detection of embedded figures is the manifestation of the fixed differential power of sensory processing mechanisms in ASD, with feedforward processes dominating perceptual processes processes (Mottron et al., 2006), demonstrated, for example, by heightened levels of brain activation during visual processing (Samson et al., 2012). Over-responsiveness to local stimuli will disrupt the integration of sensory information into perceptual wholes (percepts) and could have downstream consequences not only for low-level perceptual tasks but also for higher-level socio-cognitive tasks such as face-processing (Behrmann et al., 2006) or language processing, which depends upon the accurate and timely perception of auditory input (Kargas et al., 2015). It could also be associated with a more wide-ranging disruption of social functioning such as that outlined in Markram’s Intense World Theory (Markram & Markram, 2010).

Disruption to the information integration mechanisms underpinning normal sensory-perceptual processing (see Section 5) formed the basis of an early dysfunctional connectivity model of ASD (Brock et al., 2002) called the ‘temporal binding’ hypothesis. This argued that the evidence of atypical local and global processing in ASD (Happé & Frith, 2006; Mottron et al., 2006) might be linked to a failure in the integration of sensory information at the cortical level, caused by a reduction in the connectivity between specialised local neural networks in the brain and possibly associated with overconnectivity within isolated individual neural assemblies (Rinaldi et al., 2008). As the process of information integration or ‘temporal binding’ had been shown to be indexed by synchronised GBA (Rodriguez et al., 1999; Tallon-Baudry & Bertrand, 1999), it was hypothesised that task-specific abnormalities in GBA would be found in ASD individuals, which could characterise the condition at the cortical level and which would inform modelling of atypical connectivity in the autistic brain (Brock et al., 2002; Rippon et al., 2007).

This temporal binding model of local hyperactivity links to the neurophysiological excitation-inhibition imbalance model outlined above with, here, a focus on an imbalance caused by increased excitatory activity. It also links to translational models of autism identifying dysfunctional GABA-ergic mechanisms (Coghlan et al., 2012) and is consistent with the high incidence of epilepsy in ASD together with evidence of high levels of epileptiform cortical activity (Berg & Plioplys, 2012; Spence & Schneider, 2009).

Additionally, it links well with the ASD symptom profile, as imbalance in localised excitatory-inhibitory processes can result in anomalous perception, such as the lack of context modulation characteristic of some forms of ASD symptom patterns (Coghlan et al., 2012; Rubenstein & Merzenich, 2003; Snijders, Milivojevic, & Kemner, 2013).

The additional aspect of the model, that the localised hyper-connectivity would be associated with global hypo-connectivity (Belmonte et al., 2004; Brock et al., 2002; Casanova et al., 2002) is consistent with the idea outlined in Section 6 that autism can be characterised as a disorder of atypical brain connectivity, with global hypo-connectivity functionally linked to various spectral bands (Just et al., 2007, 2004; Khan et al., 2013) coupled with local dysregulation primarily linked to GBA and PAC anomalies. Furthermore, the model is consistent with the weak central coherence account of autism (Happé & Frith, 2006), and with the proposed feedforward/feedback processing imbalance predicted by the enhanced perceptual functioning account (Mottron et al., 2006).

### 4.2 Deficient Predictive Coding

It has also been suggested that sensory processing in ASD can be viewed within a predictive coding framework of the brain (Lawson, Rees, & Friston, 2014; Pellicano & Burr, 2012). As outlined in Section 2.2, this relates to the idea that the perceptual system makes predictions or hypotheses about the nature of upcoming stimuli, which then become matched with incoming sensory information (Friston, 2005, 2008). It has been argued that in autism these prior predictions may be less precise (Pellicano & Burr, 2012) or deployed in an inflexible manner possibly even resulting in hyper-precision (Lawson et al., 2014; Van de Cruys et al., 2013), meaning overall that perception becomes more sensitive to incoming stimuli but can be less influenced by context. This lack of top-down prediction in ASD has been suggested to underlie problems during the perception of visually ambiguous stimuli like illusory figures and could even extend to social stimuli such as facial expressions and biological movement (Lawson et al., 2014). Poor predictive coding in ASD could also render any input as apparently novel and potentially overwhelming for the system; whilst inflexible or ‘over-exact’ coding could undermine the adaptive function of ‘approximation’, allowing the incorporation of irrelevant mismatches into the anticipatory predictive code, and render an individual intolerant of novelty and change (Markram et al, 2007; Gepner and Feron, 2009; Gomot & Wicker, 2012; Lawson et al., 2014; Sinha et al., 2014; Van de Cruys et al., 2014).

As described in Section 2.2, predictive coding is linked to specific brain oscillations with a focus on gamma’s role in coding prediction errors and cross-frequency coupling (CFC) underpinning the feedforward-feedback integration processes (Voytek et al., 2010), while low frequencies mediating top-down signalling of predictions (Arnal & Giraud, 2012). Thus, as with the temporal binding approach, understanding deficient predictive coding in ASD focusses on atypical GBA but also on measures of CFC and long range connectivity in low frequencies.

Consequent on the emerging synergies between the neurocognitive and neurophysiological models of ASD, a focus of attention in neurocognitive ASD research over the last decade or so has been on “oscillopathies” (Edgar et al., 2015) and atypical connectivity patterns (Gepner & Féron, 2009). These are summarised below.

## 5. Oscillopathies and atypical connectivity in ASD

As outlined above, theories of the role of GBA (at or around 40 Hz) in ‘temporal binding’ or the formation of the coherent percepts essential for accurate information processing indicated that gamma could be a useful ‘candidate’ frequency for characterising the cortical correlates of sensory and perceptual atypicalities (Brock et al., 2002). Much of the early research into atypical cortical activity and cortical connectivity therefore focussed on gamma. It should be noted that these earlier studies therefore use a frequency range which would now be classified as ‘low gamma’ (30-60 Hz) and studied GBA predominantly in relation to its amplitude and power. Additionally, findings are based on EEG sensor-level analysis and generally focus on task-related regions of interest rather than whole-brain measures. This task-related focus together with the emphasis on temporal binding also led to comparisons between *evoked* power, where the power changes are phase-locked to the eliciting event and *induced* power, where changes are associated with but not phase-locked to the eliciting event and will show trial by trial variations. As shown in (Tallon-Baudry & Bertrand, 1999) this distinction is key where GBA is being used as a potential measure of temporal binding, as evoked changes will occur to any stimulus presentation whereas only induced power will distinguish coherent percepts, indicating the ongoing synchronising process.

### 5.1 Task-related GBA in ASD: Visual and auditory processing

Early gamma studies in autism focussed on visual processing where ASD anomalies are well documented (Dakin & Frith, 2005). Grice et al., (2001) and Brown et al., (2005) reported failures in task-related GBA to distinguish between different types of stimuli (see Table 1), upright and inverted faces in the former and presence or absence of illusory triangles in the latter. Both studies interpreted their findings in terms of the atypical GBA indexing anomalous temporal binding of features into a cohesive percept, with Brown et al., (2005) additionally noting that the high levels of GBA increases post-stimulus were consistent with deficits in neuronal excitation-inhibition balance. The localised nature of the increased GBA was also consistent with the hypothesis of increased connectivity within *local* networks (Brock et al., 2002), possibly underpinning the perceptual hyper-abilities identified as characteristic of some ASD individuals.

A study by Sun et al., (2012) provides a good example of emerging analytical possibilities in the area, with the added benefit of using MEG. Using the perception of Mooney faces, a task reliably associated with gamma generation (Rodriguez et al., 1999), they examined whole-brain measures of power and inter-trial coherence (phase-locking) at both the sensor and the source level and also considered GBA in terms of both low (25-60) and high (60-120) gamma. The ASD participants were high functioning adults. Behaviourally, the ASD group, performed worse than the control group, with longer reaction times and fewer correct identifications. Sensor level analyses revealed an *increase* in both low and high gamma power over parieto-occipital channels in control group as compared to a *decrease* in the ASD group. The control group also showed a reduction in low gamma power over frontal areas, consistent with the earlier study by Grice et al., (2001). Inter-trial coherence measures revealed reduced coherence in the ASD group over occipito-parietal areas, with greater inter-groups differences in the lower frequency band. Correlations between behavioural measures and source power in the higher frequency range revealed a different pattern in controls and in the ASD group, with only the latter showing an association between faster responses and increased GBA in atypical areas – i.e. more posterior to the typical face processing network.

A more recent study by Peiker et al., (2015) also highlights the benefits of employing coherence-based metrics in the study of GBA. Participants were required to identify moving objects presented through a narrow slit – a task requiring the integration of perceptual information across time. In both the ASD and control groups, the stimuli elicited an increase in gamma-band (40-80Hz) power. The appeal of this paradigm is that objects presented through a horizontal, but not vertical, slit requires information to be integrated across hemispheres. Indeed, Peiker et al., (2015) reported widespread gamma-band coherence in occipital cortex for the control participants, consistent with the idea that information is being passed between hemispheres (Fries, 2005, 2015). As expected, behavioural accuracy along with gamma-band coherence within the posterior superior temporal sulcus were reduced in ASD sample, specifically for the horizontal slit condition, consistent with weak central coherence accounts of the condition (Happé & Frith, 2006).

More generally, the studies of Sun et al., (2012) and Peiker et al., (2015) demonstrate the range and complexity of potential insights that can be generated from the study of task-related GBA, but also the source of possible contradictions between studies due to a variety of GBA measures. These early studies provide some support for the suggestion that anomalous ASD perception would be associated with atypical task-related gamma activity, but are limited by rather basic measures of GBA and simplistic measures of connectivity. The findings of higher or lower levels of gamma power and gamma phase in-consistency are compatible with suggestions of excitation-inhibition imbalance (Brown et al., 2005; Zikopoulos & Barbas, 2013), but clearly need a more fine-grained analysis of GBA (See Section 2) to tie it to physiological and functional aberrations. A summary of the key studies examining visual processing in ASD has been provided in Table 1.

In contrast, the study of GBA during auditory processing in ASD has benefitted from a fuller range of GBA measures. Abnormal auditory reactivity, both hypo- and hyper- reactivity, has been observed in ASD (Hazen et al., 2014; Leekam et al., 2007) and have been linked with characteristic communication difficulties (Jeste & Nelson, 2009). Orekhova et al., (2008), for instance, examined auditory sensory gating in high and low functioning ASD children by comparing the P50 ERPs to click pairs. Normal sensory gating is associated with a significant reduction in the P50 response to the 2^nd^ click; however, this reduction was significantly decreased (absent) in the low-functioning group. The ASD groups also had higher levels of gamma power and a relationship between gamma power and poor sensory gating was demonstrated, with higher gamma power correlating significantly with small or absent P50 suppression, varying as a function of the degree of impairment - with the low functioning group showing little or no P50 suppression. This atypical sensory gating in the ASD groups associated with abnormal GBA was interpreted as potentially indicating a deficit in central inhibitory circuits (Rubenstein & Merzenich, 2003; Zikopoulos & Barbas, 2013; Whittington et al., 2011; Clementz, Blumenfeld, & Cobb, 1997). The association with degree of impairment suggests that the excitation/inhibition balance is more marked in the more severely impaired children, an additional factor to consider in tracking the role of gamma abnormalities in ASD (Whittington et al., 2011).

In a more direct assessment of auditory GBA, Wilson et al., (2007), using MEG, measured steady-state responses to 500ms, monaural click trains, amplitude-modulated at 40 cycles/second to elicit a steady-state response of increased 40 Hz power in the contralateral hemisphere. The controls and the ASD group showed similar patterns of responsivity in the right hemisphere, but the ASD group showed reduced left hemisphere power with no clear 40 Hz steady state response. Gandal et al., (2010) measured gamma (30-50 Hz) power and synchronisation in the same participants, with phase-locking factor (PLF) as a measure of inter-trial coherence. There was no significant difference in induced or evoked gamma power between the ASD and the control groups, but the ASD group showed significantly reduced gamma phase-locking. A follow-up study by Edgar et al., (2015) showed higher levels of pre-stimulus power in *all* frequencies in a group of children with ASD, with smaller early evoked gamma activity to all stimuli, and decreased left and right hemisphere inter-trial coherence in the gamma band. The pre-stimulus abnormalities were most predictive of post-stimulus abnormalities and of clinical symptoms. Finally, Rojas et al., (2008) reported that in response to a monaural, 200 msec 1 khz sine-wave stimulus children with autism and their parents revealed lower evoked but higher induced gamma power and significantly lower PLF, consistent with a deficit in GBA timing and organisation. A summary of the key studies examining auditory processing in ASD has been provided in Table 2.

Across visual and auditory paradigms the variations in GBA power effects (hypo- or hyperactivity) in ASD can be reconciled with the more consistent reports of reduced phase-locking in ASD and ASD-related populations where this was also assessed, noting that reduced phase-locking will be associated with an imbalance between evoked and induced power (Rojas et al., 2011). Overall it can be concluded that high frequency brain responses in ASD consistently reveal a lack of functional organisation at the local level, hinting at suboptimal signal-to-noise ratios, likely due to an excitation/inhibition imbalance, and at deficient integration between top-down and bottom-up signals for effective perceptual processing.

**Table 1.** A summary of key electrophysiological studies into visual processing in ASD.

| Study | Modality | Participant Number | Participant Mean Age | Stimuli | Main Findings |
| --- | --- | --- | --- | --- | --- |
| Grice et al., (2001) | EEG | 8 ASD; 8 control | 36.3 ASD; 30.9 control | Upright/inverted faces | No upright/inversion difference in frontal induced gamma-band power |
| Brown et al., (2005) | EEG | 12 ASD; 12 MLD* | 14.7 ASD; 14.0 MLD | Kanisza illusory shapes | Higher induced gamma-band power both in the presence *and* absence of an illusory shape |
| Isler et al., (2010) | EEG | 9 ASD; 11 control | 7.8 ASD; 6.7 control | White light stroboscopic flashes | Earlier & greater early responsivity in alpha/beta bands; but less interhemispheric gamma-band coherence |
| Wright et al., (2012) | MEG | 13 ASD; 13 control | 15.1 ASD; 15.7 control | Emotional faces | Lower induced gamma activity in occipital areas for emotional face stimuli |
| Sun et al., (2012) | MEG | 13 ASD; 16 control | 30.3 ASD; 29.7 control | Illusory faces | Reduced inter-trial coherence in the gamma-band; mixture of regionalised increased *and* decreased gamma-band power differences |
| Stroganova et al.,<br>(2012) | EEG | 23 ASD; 23<br>control | 5.0 ASD; 5.1<br>control | Illusory figures | Weaker phase-locked beta/gamma band<br>responses, 120-270ms post-stimulus onset |
| Snijders et al.,<br>(2013) | EEG | 12 ASD; 12<br>control | 22.0 ASD;<br>22.0 control | Orientation<br>discrimination task<br>using gabor patches | Lower gamma-band power; no contextual<br>modulation effect in the gamma-band |
| Peiker et al.,<br>(2015a) | MEG | 20 ASD; 20<br>control | 31.2 ASD;<br>31.5 control | Images viewed<br>through<br>vertical/horizontal slit | Reduced occipital beta-band power; and<br>decreased gamma-band coherence between<br>bilateral superior temporal sulci |
| (Peiker et al.,<br>2015b) | MEG | 13 ASD; 14<br>control | 32 ASD; 32.1<br>control | Motion discrimination<br>task using dot<br>kinematograms | Greater gamma-band power with increasing<br>motion intensity in V3, V6 and V5 (MT) |

\*MLD = Moderate Learning Difficulty

**Table 2.** A summary of key electrophysiological studies into auditory processing in ASD.

| Study | Modality | Participant Number | Participant Mean Age | Stimuli | Main Findings |
| --- | --- | --- | --- | --- | --- |
| Orekeva et al., (2008) | EEG | 21 ASD; 21 control | 5.9 ASD; 6.0 control | Paired clicks | Higher gamma power; P50 suppression reduced for the severely impaired ASD subjects |
| Wilson et al., (2007) | MEG | 10 ASD; 10 control | 12.4 ASD; 12.0 control | Monoaural clicktrain | Reduced 40Hz steady state response in left auditory cortex |
| Gandal et al., (2010) | MEG | 25 ASD; 17 control | 10.2 ASD; 10.7 control | Sinusoidal tones | 10% delay in the M100 evoked response; and a reduction in phase-locking for the gamma band |
| Edgar et al., (2015) | MEG | 105 ASD; 36 control | 10.1 ASD; 10.1 control | Sinusoidal tones | Elevated pre-stimulus power across from 4-80Hz; smaller evoked (50-150ms) gamma response; decreased inter-trial coherence in the gamma-band |
| Rojas et al., (2008) | MEG | 11 ASD; 16 control | 31.5 ASD; 43.1 control | Sinusoidal tones | Reduced evoked gamma; increased induced gamma; reduced phase locking in the gamma-band |
| Roberts et al.,<br>(2010) | MEG | 25 ASD; 17 control | 10.8 ASD;<br>10.2 control | Sinusoidal tones | Delay in M100 evoked response |
| (Port et al., 2016) | MEG | 27 ASD; 9 controls<br>(longitudinal) | 8.4/12.4<br>ASD;<br>8.1/12.1<br>control | Sinusoidal tones | Delay in M100 evoked response; reduced<br>gamma-band evoked activity at both time<br>points |

### 5.2 Task –related cross-frequency, phase-amplitude coupling (PAC) in ASD

Where the interest in gamma is as a measure of cortical connectivity, the recent use of phase amplitude coupling (PAC) metrics has provided further insights into ASD (See Section 2). Given that theoretical models of autism have focussed on information integration (Rippon et al., 2007; Vissers et al., 2012), measures of PAC between gamma and lower frequency oscillations could prove very informative (e.g. Voytek & Knight, 2015), especially in the context of local to global brain connectivity in ASD. Indeed, Khan et al., (2013) have recently employed a PAC metric to index local connectivity in relation to face stimuli in the fusiform face area (FFA). In their MEG study the authors examined gamma power and alpha-gamma coupling in the fusiform face area in young male ASD participants and matched controls in response to neutral or emotional faces as compared to houses. There were no group differences in evoked responses in either the alpha or gamma band. Long-range connectivity was measured using broadband (6-55 Hz) coherence and revealed lower levels in the ASD group. Alpha-gamma coupling measures revealed reductions in local functional connectivity in the fusiform face area in ASD, with the gamma effects in the high frequency range (75 -110 Hz). It is important to note that these PAC differences emerged despite the failure of both alpha and gamma power measures alone to distinguish between the groups and implicates the timings of any gamma related changes rather than the power per se as potential distinguishing features. Additionally, the PAC measures were shown to be negatively correlated with ADOS scores, thus providing a useful biomarker for symptom severity, and also, using classifier techniques, successfully distinguishing the ASD and control group with 90% accuracy.

In auditory/speech models it has been shown that auditory cortical responses to speech occur in the theta range, which then modulates gamma activity (Giraud & Poeppel, 2012; Schroeder et al., 2008). Jochaut et al., (2015), using fMRI and EEG data, examined theta (4-7 Hz)-gamma (30-40 Hz) coupling in response to continuous speech in a heterogeneous group of ASD participants with IQ scores ranging from 35-124, and including dysphasic as well as linguistically normal participants. They noted that there were significantly higher pre-stimulus theta levels in the auditory cortex in the ASD group which did not increase with speech stimulation. Subsequent examination of theta-gamma relationships demonstrated that theta activity in the left auditory cortex did not vary as a function of speech modulations and failed to down-regulate gamma oscillations in the ASD group. This would be equivalent to the anomalous gamma activity described in the preceding studies, particularly those that consistently reported lacking phase relationships in gamma as well as higher levels of induced but not evoked gamma. Additionally, the theta-gamma measure predicted verbal ability in the AD group (r=0.746, p=0.008) and was strongly tied to the general autism symptoms. Further, examining EEG-BOLD coupling allowed assessment of the connectivity between auditory cortex and speech areas and indicated reduced connectivity from A1 to Broca’s area and the motor cortex, but not the other way round, suggesting the theta-gamma anomaly is primarily sensory. This provides an explanatory model for the sensory abnormalities in ASD and also is strong support for the notion that the origins of atypical ASD behaviour may lie in more fundamental sensory and perceptual dysfunctions. A key aspect in assessing past studies and designing future ones is of the choice of gamma and gamma-related metrics; it is clear that power measures alone are not sufficiently sensitive. In addition, where local connectivity is an aspect of interpretation of GBA, measures of phase-amplitude coupling, PAC, rather than simple coherence could prove extremely useful.

Recently, computational modelling of oscillatory activity indeed suggests that PAC may be a key component in balancing excitation-inhibition interactions and maximising information flow between brain areas (Peterson & Voytek, 2015).

The proposed dysfunction of oscillations and functional connectivity, especially at the local level and in relation to GBA and PAC measures, are consistent with neurophysiological models of ASD at the cellular level. For instance, computational modelling of oscillatory activity suggests that PAC may be a key component in balancing excitation-inhibition interactions (Peterson & Voytek, 2015). Voytek & Knight (2015) therefore proposed deficient PAC as an index for a local excitation/inhibition imbalance in various psychopathologies and in ASD in particular, which could be due to reduced inhibition (Vogels & Abbott, 2009) or an affected excitation-inhibition ratio (Rubenstein & Merzenich, 2003) as discussed in Section 2.

### 5.3 Task-related and task-free global oscillatory connectivity deficits in ASD

Studies of GBA during auditory and visual processing in particular have proved to be a useful testing ground for the application of different ways of measuring task-related GBA and linking this to hypothesised differences in cortical connectivity. Findings are complex but, on the whole, support models implicating greater reactivity in the early sensory processing stages combined with an apparent failure to ‘titrate’ such reactivity as a function of the stimulus characteristics (e.g. upright vs. inverted faces, face vs houses, presence or absence of illusory figures). This inability to regulate local sensory gamma-band activity may be a reflection of atypical patterns of oscillatory activity within lower frequency bands, which are thought to co-ordinate so-called ‘top-down’ long range connectivity (Engel, Fries, & Singer, 2001; Jensen et al., 2015). Khan et al., (2013) for instance, did not only report reduced alpha-gamma PAC in relation to face processing in ASD, reflecting local hypo-connectivity and dysregulation, but also employed alpha-phase coherence as a measure for long-distance (global) connectivity, showing a reduction in this measure too. In an MEG picture-naming study looking at functional connectivity in ASD as measured by Granger causality, Buard et al., (2013) reported higher beta-and gamma band functional connectivity in the autism group than in controls. Isler et al., (2010), used a long-latency flash visual evoked potential (VEP) task in a small group of young ASD children (5.5-8.4 years old) and reported that inter-hemispheric coherence and phase synchrony were reduced in the ASD group at all frequencies, particularly those above theta.

These results are consistent with findings from task-based fMRI studies using ASD individuals, with the general consensus being reduced long-distance connectivity (e.g. (Schipul, Keller, & Just, 2011). However it remains unclear how these results relate to patterns of atypical local connectivity reported using fMRI (e.g. Itahashi et al., 2015; Keown et al., 2013), and conflicting results are often reported. For example, You et al., (2013) used fMRI to compare functional connectivity during resting state and sustained attention in ASD and control groups. Task-related distant functional connectivity maps were more focal than resting state maps in the control group, but were more diffuse in the ASD group. However no group differences were found in resting or task-based local connectivity.

This lack of consistency between findings of atypical local and global connectivity in the autistic brain may be a function of the methodology, related to the lower levels of temporal resolution in fMRI. MEG studies, however, offer greater temporal sensitivity, and generally produce a more complex pattern of results which may ultimately help to reconcile apparent contradictions within the fMRI literature. For example a recent study by (Khan et al., 2015) used 25Hz vibrotactile stimulation to study somatosensory processing in autism. The authors found increased feedforward connectivity from primary to secondary somatosensory cortex in the ASD group, accompanied by reduced phase-locking at 50Hz which the authors argued was evidence of atypical recurrent processes at the local level. This result suggests that the direction of connectivity, as well as the specific type of neural activity needs to be taken into account when studying ASD. As evidenced in this section, oscillatory measures based on coherence, granger causality and PAC offer more nuanced insights into patterns of both local and global task-based connectivity in the autistic brain (Khan et al, 2013) and may be able to reconcile previously contradictory findings in the field.

In addition to atypical sensory responsivity, consistent reports of high levels of epileptiform activity in ASD cortical activity as well as the high incidence of epilepsy (Berg & Plioplys, 2012), suggests that there could be unusual levels of high-frequency activity in resting state EEG/MEG activity in ASD populations. The study of electrophysiological resting-state activity in autism has generally been in the context of network-based inter-regional connectivity, using simple coherence-based measures or more complex metrics based on graph theory, and also in the identification of resting-state networks (Brookes et al., 2011). In ASD research, early work using MEG/EEG, reported anomalies in lower frequency bands, such as patterns of both inter- and intra-hemispheric hypoconnectivity as measured by coherence (Coben et al., 2008; Murias et al., 2007), as well as increases in high frequency power (70-120 Hz) particularly in posterior brain regions (Cornew, 2012). More recent work has focussed on localising these resting-state oscillopathies. For example, Kitzbichler et al., (2015) collected data from 15 ASD participants, aged 6-21, and mapped this to source-space using a minimum norm estimate. Using various graph theory metrics, the authors showed that in the gamma band (30-70 Hz), the ASD group showed stronger and more efficiently connected networks, with many more connections from occipital areas to parietal, temporal and frontal regions. In the beta band, the ASD group were characterised by less efficiently connected networks, particularly those involving the frontal/parietal lobes. There were also group differences in age-related connectivity changes, with their (small) ASD group showing little evidence of connectivity-based maturation, as compared to clear evidence of developmental changes in controls. Overall, the authors interpret these differences as a developmental imbalance between feedforward mechanisms, primarily mediated by GBA, and fronto–parietal regulatory feedback mechanisms, mediated by lower-frequency oscillations. This clearly links with results of dysregulated gamma-band activity within sensory regions of ASD participants (Cornew 2012) and atypical neurophysiological signatures of frontal lobe function.

Overall, resting state measures offer a fruitful approach of the study of ASD, not least because they offer the opportunity of involving younger and/or lower functioning participants. Key findings from this area are summarised in Table 3. The possibility of characterising the network connections using graph theory (Sporns, 2003; Stam & Reijneveld, 2007) and demonstrating how these can be associated with symptom patterns (Kitzbichler et al., 2015) supports the findings from studies of auditory gamma that GBA could serve as a potential biomarker for the condition; although this possibility remains speculative at the present time. However, it is also important to note that resting state studies measure intrinsic brain activity, which may be unable to elucidate the full range of connectivity differences between autistic participants and controls (Morcom & Fletcher, 2007). Future research should therefore attempt to determine the relationship between resting state and experimentally-driven measures within the same individuals (GBA, PAC, long-range connectivity, etc.).

**Table 3.** A summary of key task-free/resting-state ASD studies using EEG and/or MEG

| Study | Modality | Participant Number | Participant Mean Age | Main Findings |
| --- | --- | --- | --- | --- |
| Murias et al., (2007) | EEG | 18 ASD; 18 control | 22.7 ASD; 24.9 control | Pattern of higher *and* lower power between 3-17Hz; higher theta-band coherence in frontal regions; lower alpha-band coherence |
| Tierney et al., (2007) | EEG | 65 high-risk ASD infants; 57 low-risk infants | Developmental sample (6-24 months) | High risk infants showed lower spectral power across all frequency bands and different developmental trajectories |
| Coben et al., (2008) | EEG | 20 ASD; 20 control | 8.9 ASD; 9.0 control | Higher theta and delta power, especially in frontal electrodes; lower interhemispheric coherence; higher coherence in temporal regions |
| Sheikani et al., (2009) | EEG | 15 ASD; 11 control | 9.2 ASD; 9.1 control | Lower gamma power in frontal and temporal electrodes |
| Pollonini et al., (2010) | MEG | 8 ASD; 8 control | 18.7 ASD; 19.0 control | Graph theoretic analysis showed increased path-length |
| Cornew et al., (2012) | MEG | 27 ASD; 23 control | 9.8 ASD; 10.8 control | Higher power across multiple frequency bands, including alpha and gamma |
| Maxwell et al.,<br>(2015) | EEG | 15 ASD; 18 control | 15.1 ASD; 14.2<br>control | Lower gamma power in right lateral electrodes |
| Kitzbichler et al.,<br>(2015) | MEG | 15 ASD; 15 control | 12.5 ASD; 13.0<br>control | Graph theoretical analysis showed stronger gamma connections in occipital cortex; but reduced frontal connectivity in theta, alpha + beta frequency bands |
| Bartfield et al.,<br>(2007) | EEG | 10 ASD; 10 control | 23.8 ASD; 25.3<br>control | Graph theoretical analysis showed increased path length in the delta band; greater numbers of short-range connections but lower numbers of long-range connections |

### 5.4 Summary

This section has reviewed the past research regarding oscillopathies in ASD, with a focus on the transition from traditional-power based metrics to emerging measures of phase-based connectivity measures. Traditionally GBA measures were the focus of research, with GBA power indices revealing a rather inconsistent pattern, with reports of hypo- as well as hyper-activity (Section 5.1). In contrast, novel measures such as local inter-trial phase coherence, local PAC and global phase-to-phase coupling seem to be promising in terms of revealing deficits more reliably and potentially being able to absorb ASD idiosyncrasies. These measures may therefore complement or even replace traditional power measures, yet it remains unclear if and how these measures might relate to each other and what the pattern of identified oscillopathies might reveal about aberrant function in ASD.

PAC has been proposed as a measure of local processing integrity and the very few reports to date (reviewed in Section 5.2) indicate that there could be an ASD deficit in PAC even in the absence of a gamma power deficit (e.g. Khan et al., 2013). Long-range connectivity in the form of phase-coupling is assumed to reflect cross-systems information integration and ASD-specific abnormalities have been identified between various brain areas and in various frequencies (reviewed in Section 5.3). However, specific predictions regarding which frequencies (delta, theta, alpha, beta, gamma) and connections (e.g. top-down vs. bottom-up) are expected to reflect hyper- in contrast to hypo-connectivity and in what particular conditions (e.g. task-related vs. task-free) are only emerging and would have to be based within a systematic theoretical framework that is currently missing. Furthermore, such a framework should also allow the prediction of whether different measures of oscillopathies would have to be conceived of as independent or as mechanistically linked. For instance, deficient local PAC could be related to long-range connectivity in form of insufficient low frequency coupling across brain areas that in turn may not succeed in entraining high frequencies locally (e.g. Arnal & Giraud, 2012; Mejias et al., 2016; Onslow et al., 2014; Peterson & Voytek, 2015), which might also be reflected in deficient inter-trial phase coherence in low and high frequencies. Finally, even if such a mechanistic link was identified reliably, the necessity and the role of PAC and long-range coupling for cognitive function will have to be addressed within a consistent theoretical framework of ASD. In the following section such an explanatory framework will be proposed, linking together several measures of oscillopathies in an attempt to bridge the gap between electrophysiology, cognitive function and ASD symptomatology.

## 6. A novel approach to ASD: local dysregulation, global hypoconnectivity, and deficient predictive coding

As reviewed in Section 5, the overall pattern of findings regarding basic GBA in autism could be regarded as inconsistent, with some studies reporting higher while other reporting lower gamma power in various sub-bands (Section 5.1). However, this apparent inconsistency could be reconciled by characterising these findings as evidence of a local dysregulation of optimal processing that may present as *either* increased or decreased gamma band power depending on context and task and/or on the symptom profile of the ASD cohort. This would mirror the heterogeneity evident in ASD, a factor which is noted in other reviews of ASD research and interpretation (Dickinson et al., 2016; Picci et al., 2016; Simon & Wallace, 2016).

### 6.1 Core features of the proposed framework

The notion proposed here shifts the explanatory focus away from the question of local hyper- vs. hypo-connectivity in ASD as taken to be reflected by gamma power alone (e.g. Brock et al., 2002). Instead it suggests local dysregulation may be the underlying cause of connectivity deficits, as evidenced by deficient cross-frequency coupling PAC (e.g. Khan et al., 2013) and/or evidence of excitation/inhibition imbalance (Casanova et al., 2002; Rubenstein & Merzenich, 2003; Zikopoulos & Barbas, 2013) (see Sections 2.1, 4.3). As shown in Figure 1, PAC relies on (or even constrains, e.g. Peterson & Voytek, 2015) the local excitation/inhibition balance, thus possible future research should focus on complementary measures to corroborate this link. Another critical implication of focussing on PAC (and other CFC measures) is the strong mechanistic link between high and low frequency brain oscillations (see Figs, 1 and 2) that may lead to testable predictions. These are two of a few testable assumptions emerging from the proposed framework, which are listed in Table 4.

**Figure 2.**
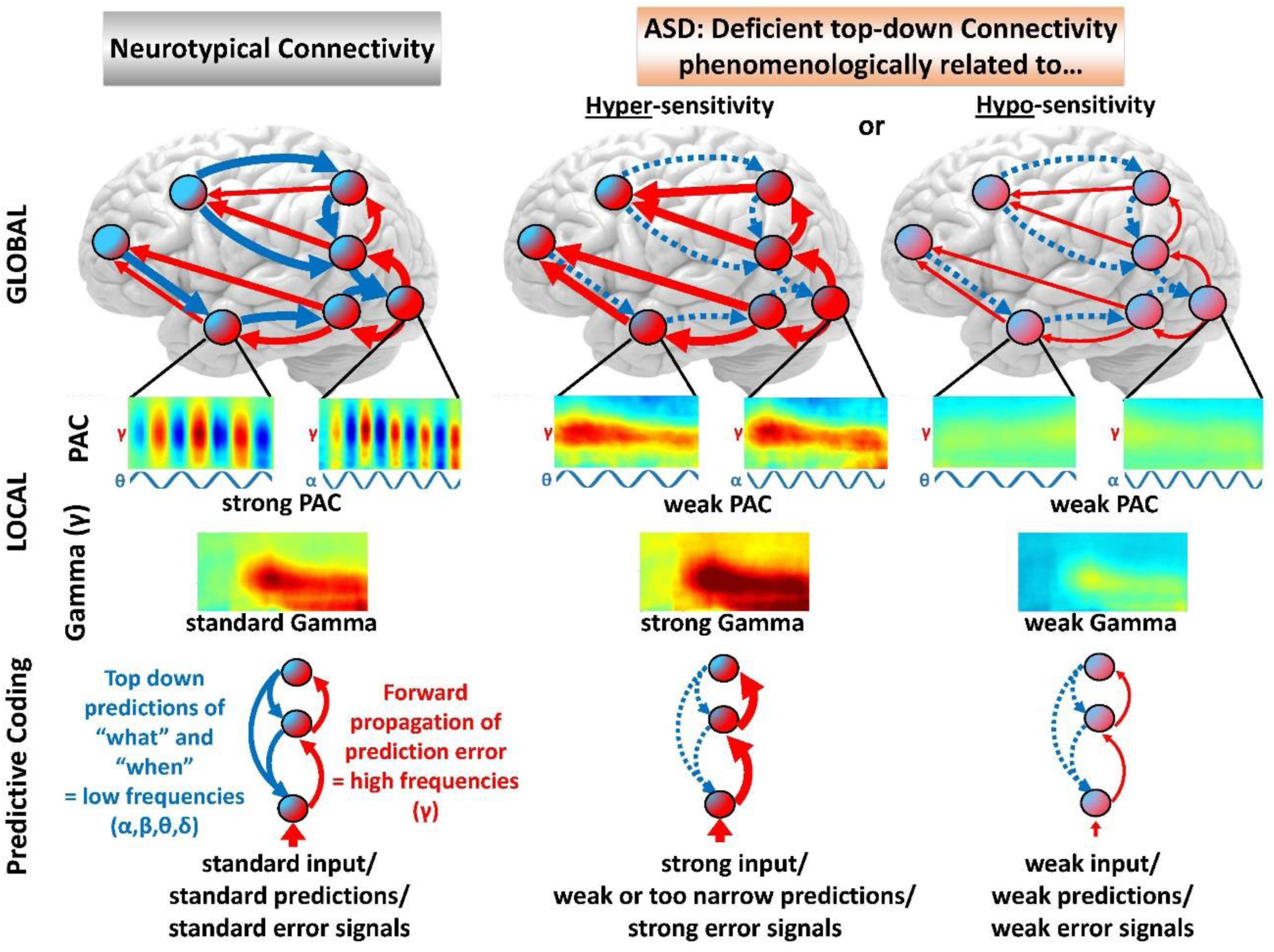
Schematic illustration of the proposed framework. The left column shows assumed neurotypical connectivity and measures, while the middle and right columns show hypothetical ASD connectivity and measures for the case of hyper-sensitivities (middle) and hypo-sensitivities (right), respectively. The top row shows schematic functional connectivity at the global level, with low frequency (delta δ, theta θ, alpha α, beta β) top-down connections in blue and high frequency (gamma γ) bottom-up connections in red. The ratio of red/blue in individual nodes reflects whether top-down or bottom-up influences are thought to prevail. Top-down, feedback connectivity in low frequencies is assumed to be deficient in both expressions of ASD sensory sensitivities (hyper- and hypo-), while bottom-up, feedforward connectivity in γ is assumed to be overexpressed for hyper-sensitivities (middle), yet underexpressed for hypo-sensitivities (right). The middle row shows measures of local processing in form of phase-amplitude-coupling (PAC) at the top and γ power underneath. Significant entrainment of local γ by low frequencies (e.g. α, θ shown here) has been reported for neurotypical participants (see text), while it is proposed here that in ASD such entrainment should be deficient (e.g. Khan et al., 2013). Note that it is further hypothesised (see also Table 4) that gamma power could be overexpressed for ASD hyper-sensitivities (middle), despite deficient PAC (γ is strong but not entrained by lower frequencies), while γ power could be underexpressed for ASD hypo-sensitivities (right). Overall deficient PAC in ASD could be related to deficient low frequency coupling at the global level: Deficient top-down coupling in α or θ should also be reflected in deficient entrainment of γ by α or θ at the local level (see Table 4). The bottom row shows predictive coding which is hypothesised to be deficient in ASD (middle and right). Based on Arnal & Giraud (2012) and in alignment with our general framework (top row) top-down predictions are assumed to encode “what” is expected and “when” via lower frequencies (δ, θ, α, β,), while γ is assumed to propagate error signals up the processing hierarchy. Again over- vs. under-expressed γ connectivity and power should be observed for hyper- (middle) and hypo-sensitivities (right), respectively.

The proposed local dysregulation (as reflected, for instance, by PAC, see Figure 2) has therefore at least three mechanistic consequences and associated functional aberrations (see Fig. 2). Firstly, as described, local functioning will be affected as it is hard to achieve optimal signal-to-noise ratios (SNR) with a suboptimal balance between excitation and inhibition (Voytek & Knight, 2015); this will be reflected by reduced PAC (Khan et al., 2013). Such an imbalance may result in ASD-typical sensory hyper-sensitivities, because strong external stimuli generate strong incoming signals via pyramidal neurons that may result in excessive local activation if neural structures for effecting suppression are underdeveloped (e.g. Rubenstein & Merzenich, 2003). However, while ASD individuals commonly report hyper-sensitivity to arousing stimuli, hypo-sensitivity is also reported for a subset of individuals (Ben-Sasson et al., 2009). We therefore propose that hyper-sensitivities and hypo-sensitivities may not necessarily reflect local hyper- or hypo-connectivity, but rather a fundamental imbalance between excitation and inhibition that may be expressed differently in different individuals and possibly in relation to different stimuli. At the same time such an imbalance may also explain “confusion” in signal processing when several features or stimuli are equally strong or weak (SNR is too low), resulting in abnormally alternating and /or incoherent stimulus representations (e.g. merged binocular images and longer switch times reported by Robertson et al., (2013) that may further explain the subjective impression of many autistic individuals that the “world is too intense” (Markram & Markram, 2010), as well as the tendency to neglect global gestalts (Bölte et al., 2007; Scherf et al., 2008; Walter et al., 2009), as discussed in Sections 1 and 3, that would require more systematic coordination of local processing. For dissociating hypo- and hyper-sensitivities, it is proposed (see Figure 2, middle vs. right column) that hypo-sensitivities in ASD should be reflected by *low* gamma power, weak PAC and inter-trial phase coherence, as well as by deficient top-down connectivity in low frequencies. In contrast, hyper-sensitivities in ASD should be reflected by *strong* gamma power, but weak PAC and inter-trial phase coherence, as well as by deficient top-down connectivity in low frequencies (Table 4, Hypothesis 3).

Secondly, in agreement with an argument initially proposed by Rubenstein & Merzenich (2003) long-range connectivity will be affected if local processing is not reliable. Developmentally this may impact on the establishment of cross-cortical connections, especially resulting in deficient links with higher-level control areas such as the prefrontal cortex (e.g. McPartland & Jeste, 2015). While gamma feed-forward connectivity could even be overexpressed (Kitzbichler et al., 2015) due to the described excitation/inhibition imbalance at the local level, establishment of reciprocal top-down connectivity associated with lower frequency bands (Arnal & Giraud, 2012; Cavanagh & Frank, 2014; Kitzbichler et al., 2015) would be hampered due to the rather erratic and unsystematic nature of the local sensory processing. However, it is equally conceivable that a lack of top-down long-range connectivity (from prefrontal cortex; e.g. Kitzbichler et al., 2015; Schipul et al., 2011); to sensory areas lies at the core of the problem, resulting in a lack of top-down regulation of the local excitation/inhibition balance. Application of directional measures of long-range connectivity such as Granger causality (e.g. Michalareas et al., 2016) and/or dynamic causal modelling (Friston, Moran, & Seth, 2013; Penny et al., 2009) to MEG/EEG task data collected from ASD participants may be able to elucidate this complex interplay between local and global processing levels.

Whatever the exact primary cause, the result will be further dysregulation at the local processing level due to a lack of top-down influence on signal selection, filtering, and integration, accompanied by a local imbalance in excitation vs. inhibition resulting in deficient SNR: either too high, resulting in hypersensitivity (e.g. high GBA, PAC and/or inter-trial coherence), or too low, resulting in erratic/incoherent processing (e.g. low GBA, PAC and/or inter-trial coherence). Visual sensory dysfunctions in ASD may serve as a theoretical example of the affected interplay between bottom-up and top-down processing. In concordance with the framework proposed here (Figure 2), Michalareas et al., (2016) have recently reported a connectivity hierarchy of the neurotypical visual system based on Grainger causality measures of frequency-specific connectivity that revealed a predominance of feedforward, bottom-up connectivity in gamma frequencies, yet a predominance of feedback, top-down connections in the alpha/beta range. Figure 3 summarises these findings and extrapolates to the case of ASD (see Table 4, Hypothesis 4): According to the proposed framework (also Fig. 2) top-down, feedback connections are affected in ASD, thus, the effects reported by Michalareas et al., (2016) for alpha/beta should be absent in ASD, i.e., no discernible advantage for feedback connections in alpha/beta should be observed. In contrast, gamma feedforward connectivity might be overexpressed in individuals with visual hypersensitivities (yet under-expressed for individuals with the more infrequent hypo-sensitivities).

**Figure 3.**
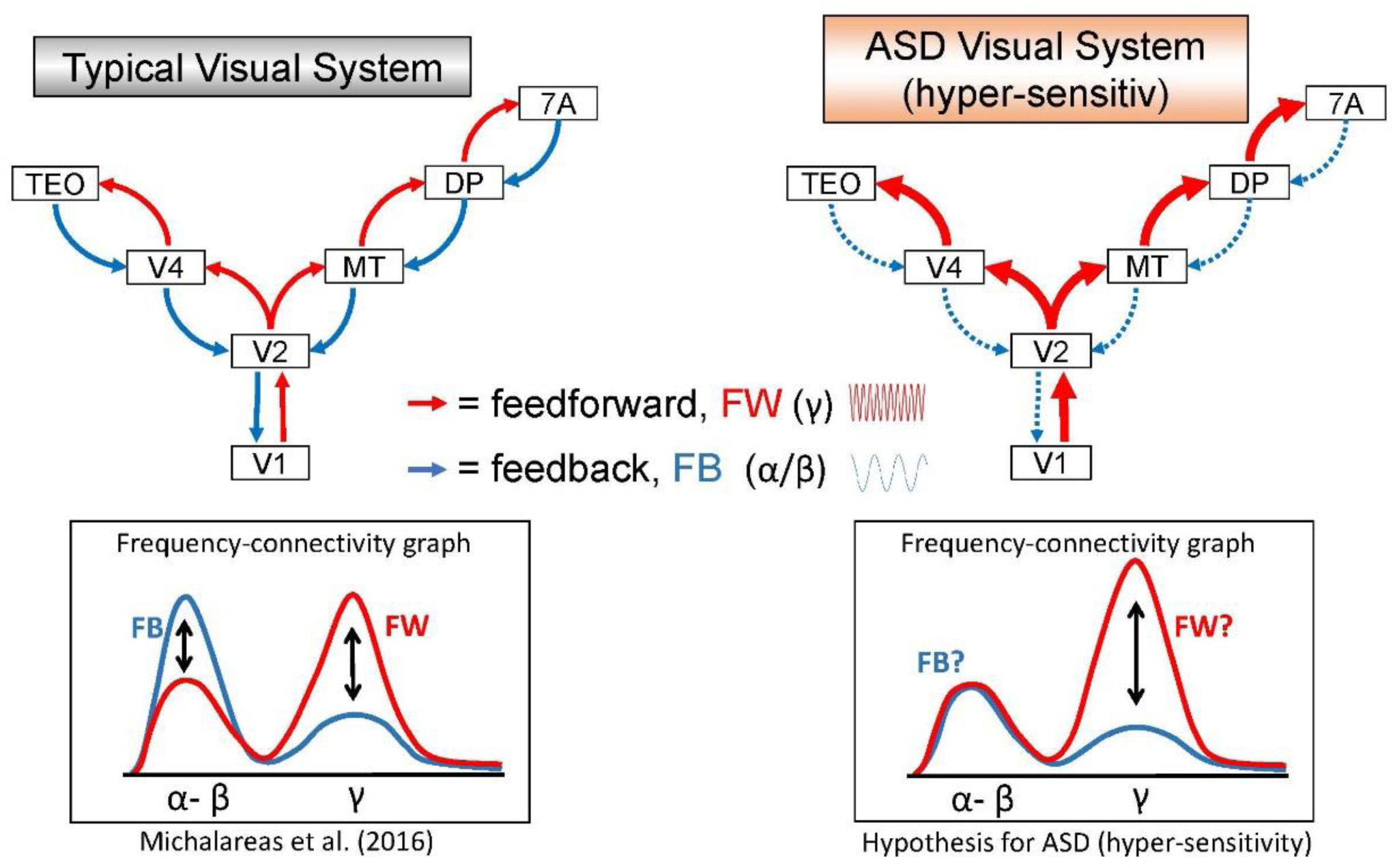
Diagram to illustrate our proposed framework for atypical early visual processing in ASD (for simplicity only the case of hyper-sensitivities is shown on the right, but can be extrapolated to hypo-sensitivities in concordance with Fig. 2). Seven putative regions of the visual system are shown that have known neurotypical feedforward/feedback connectivity profiles as described by Michalareas et al. (2016). The frequency-connectivity plot at the bottom left (adapted from Michalareas et al., 2016) reveals that feedback connectivity is dominant in the alpha/beta range, while feedforward connectivity prevails in the gamma range. The frequency-connectivity plot at the bottom right predicts that in ASD the feedback connectivity advantage for alpha/beta would not be observed, potentially in conjunction with a more pronounced feedforward effect for gamma (-in case of hyper-sensitivities; for hypo-sensitivities generally attenuated gamma might be observed).

The combined effects of local dysregulation and deficient top-down control discussed above, lead to the third proposed consequence. An optimal balance in SNR including top-down regulation in form of selection and filtering allows for optimal predictive coding of the environment (see also Lawson et al., 2014; Pellicano & Burr, 2012; Van de Cruys et al., 2013). With intact top-down regulation (filtering, selection, integration) local encoding can be predictive for “what” should happen and “when” (Arnal & Giraud, 2012), thus mainly processing deviations from expectations, resulting in a system that is highly efficient and proactive to respond (see Fig. 2, bottom row; Fig. 3). However, if top-down regulation is absent (due to a lack of long-range connections or a lack of an effect on locally dysregulated circuits), then the system could be forced away (developmentally) from predictive coding to a purely reactive process, with the “world” becoming progressively unpredictable without (therapeutic) intervention and the system tending to shut itself off from “erratic” input and/or seeking “sameness and reassurance” in self-initiated repetitive behaviours that increases predictive success while reducing the amount of prediction error (Kargas et al., 2015; Lawson et al., 2014; Van de Cruys et al., 2013).This would be consistent with the Bayesian explanations of autistic perception as described in Section 4.2 (Pellicano & Burr, 2012). In fact, the framework proposed here is more aligned with Pellicano & Burr's (2012) notion of “weak priors”, where predictions about the world remain underspecified (hypo-precise), thus, external input to the system is perceived as “surprising” in the absence of adequate predictions, either generating aberrant error signals and/or unmodulated bottom-up stimulus processing as a consequence (see Fig. 2, bottom row). In contrast, accounts that propose “hyper-precision” (Lawson et al., 2014; Van de Cruys et al., 2013), suggest that predictions in ASD about the world are so narrow and precise that external input to the system is very likely to deviate, thus, also generating strong aberrant error signals. Within the framework proposed here a testable hypothesis can be formulated that distinguishes between the two accounts (Table 4, Hypothesis 5): While both accounts would hypothesise strong gamma feedforward connectivity due to either predominantly bottom-up processing or strong deviant error signals (conforming to Arnal & Giraud, 2012), the hypo-precision account predicts *deficient* top-down connectivity in lower frequencies that is supposed to feedback predictions to lower processing levels (see Figures, 2, 3, 4), while the hyper-precision account predicts *typical or stronger* top-down connectivity in lower frequencies that feeds back the hyper-precise expectations about the external world (conforming to Arnal & Giraud, 2012).

**Table 4.** A list of hypotheses associated with our novel framework for ASD

|  |
| --- |
| **Hypothesis 1 (Sections 2.2, 6.2, Figure 1):** |
| PAC is proposed to be tightly linked with local excitation/inhibition balance (Fig. 1), thus future research should focus on complementary measures to corroborate this link in typical and ASD participants (e.g. correlations between PAC and GABA). |
| **Hypothesis 2 (Sections 2.3, 6.2, Figures 1, 2):** |
| A critical implication of focussing on PAC (and other CFC measures) is the strong mechanistic link between high and low frequency brain oscillations (see Figs, 1 and 2) that should be reflected by PAC in conjunction with low frequency cross-cortical coupling in typical participants *while ASD participants should show <u>deficient</u> PAC and associated long-range low frequency coupling.* |
| **Hypothesis 3 (Section 6.2, Figure 2):** |
| It is predicted that <u>hypo</u>-sensitivities in ASD should be reflected by *low gamma power*, weak PAC and inter-trial phase coherence, as well as by deficient top-down connectivity in low frequencies. In contrast, <u>hyper</u>-sensitivities in ASD should be reflected by *strong gamma power*, but weak PAC and inter-trial phase coherence, as well as by deficient top-down connectivity in low frequencies. |
| **Hypothesis 4 (Section 6.2, Figure 3):** |
| According to the proposed framework (also Fig. 2) top-down, feedback connections are affected in ASD, thus, the effects reported by Michalareas et al (2016) for *alpha/beta feedback connectivity should be <u>absent</u> in ASD*, i.e., no discernible advantage for feedback connections in alpha/beta should be observed. In contrast, *gamma feedforward connectivity might be overexpressed* in individuals with visual hypersensitivities yet potentially under-expressed for individuals with the more infrequent hypo-sensitivities. |
| **Hypothesis 5 (Section 6.2):** |
| While both types of predictive coding accounts of ASD (<u>hypo</u>- vs. <u>hyper</u>-precision) would hypothesise strong gamma feedforward connectivity (based on the framework proposed here), the <u>hypo</u>-precision account predicts *deficient top-down connectivity in lower frequencies* that is supposed to feedback predictions to lower processing levels (see Figures, 2, 3, 4), while the <u>hyper</u>-precision account predicts *typical or stronger top-down connectivity in lower frequencies* that feeds back the hyper-precise expectations about the external world. |
| **Hypothesis 6 (Section 6.3):** |
| It is predicted that functional gamma feedforward connectivity (reflecting aberrant error signals) as well as aberrant low frequency feedback connectivity (reflecting deficient top-down predictions) *should normalise in response to external feedback and during repetitive behaviours compared to novel actions.* |
| **Hypothesis 7 (Section 6.3, Figure 4):** |
| Deficiencies in dynamic decoding of social stimuli and sequences (temporal integration required) should be primarily related to low frequency aberrations (delta, theta) and associated PAC, while more static stimuli could reveal deficient PAC in relation to higher alpha/beta frequencies. |
| **Hypothesis 8 (Section 6.3, Figure 4):** |
| We hypothesise that especially *low delta-theta frequency long-range phase coupling should be affected* in ASD in conjunction with *reduced local PAC and possibly inter-trial phase coherence* during high-level social cognition that requires complex signal integration over time. |

### 6.2 Explaining ASD symptomatology beyond sensory aberrations

It is important to point out that the current framework does not aim at explaining all ASD symptoms in full. In contrast to various other approaches, however, we have considered idiosyncrasies in ASD to some extent, e.g. by proposing that local dysregulation and deficient top-down input may be at the root of hyper- as well hypo-sensitivities reported for different individuals with ASD (Figs. 2, 3, 4). The proposed framework provides a consistent platform based on common electrophysiological mechanisms that assumes similar oscillopathies throughout the cortical system, which may manifest as various functional aberrations depending on the involved subsystems. The implications for sensory aberrations of our predictive account based on local dysregulation and deficient top-down connectivity have been discussed in detail in the previous section, and could be extended to motoric aberrations and social symptoms as well.

This proposed framework can also be applied to the study of motor control in autism, with a focus on predictive-coding, forward and inverse models of action control (Shipp, Adams, & Friston, 2013). Motor control deficits have been consistently reported in ASD and have been related to deficient forward modelling of sensorimotor outcomes (Lawson et al, 2014; Pellicano & Burr, 2012, for reviews). In other words, the ASD system is less effective at predicting what the effects of its motor actions will be, i.e., how the world will have changed based on the action and, importantly, how the body will “feel like “while performing the action. Corroborating evidence comes from feedback training with ASD participants, where visual feedback about their body posture benefits their performance (Somogyi et al., 2016), supporting the notion of affected sensorimotor predictions and feedback loops that can be strengthened via added external feedback. Neurophysiologically, this may be linked to reports of cerebellum dysfunction in autism and associated impairments to long-distance cerebello-thalamo-cortical connectivity (Fatemi et al., 2012), meaning that the motor system is unable support the complex predictive coding required for intricate motor tasks (Wang, Kloth, & Badura, 2014). Another relevant aspect in the context of a world that is hard to predict by the ASD system is the frequent occurrence of repetitive actions and behaviours in ASD. As described above this is an expected outcome for a system where predictions are either too weak or too narrow to make coherent sense of sensory input: Repeating the same action again and again engages a loop that is much more predictable than the rest of the world, possibly reducing the bombardment with unmodulated sensory input and/or constant error signals within the system. This leads to the testable hypotheses (in agreement with

Arnal & Giraud, 2012, see Table 4, Hypothesis 6) that functional gamma feedforward connectivity (reflecting aberrant error signals) as well as aberrant low frequency feedback connectivity (reflecting deficient top-down predictions) should normalise in response to external feedback and during repetitive behaviours compared to novel actions.

Within the proposed predictive framework social and non-social stimuli and scenarios would differ primarily on the dimension of their predictiveness (see also Lawson et al., 2014) and the general hypothesis would be that ASD participants would be generally hampered in generating adequate predictions, particularly so with respect to stimuli that require complex, context-specific predictions under high uncertainty and ambiguity. Unsurprisingly, substantial parts of the cortex are dedicated to processing social stimuli, decoding and predicting others’ expressions, actions, intentions, and communication as well as planning own responses and actions (see Figure 4, Panel A). Accordingly, social stimuli such as faces and bodies seem to require more complex perceptual predictions (or “priors” within the Bayesian framework) than standard non-social objects, which is typically subsumed under the label of “holistic” processing. The existence of such specific priors that predict the stimulus as a whole seems to be corroborated by inversion effects, where efficiency of predictions collapses when a face or a body is presented upside down (Reed et al., 2003; Valentine, 1988). ASD participants have been reported to have reduced face inversion effects conforming to a strategy that is generally biased towards local feature processing compared to more holistic processing in neurotypical participants (Reed et al., 2003; Valentine, 1988) ASD participants have been reported to have reduced face inversion effects conforming to a strategy that is generally biased towards local feature processing compared to more holistic processing in neurotypical participants. Thus, the notion presented here is in agreement with these observations and with the proposal that deficient global or holistic processing in general may be at the root of various social as well as non-social perceptual aberrations in ASD (Lawson, et al., 2014).

While social deficits in ASD have been observed using static face stimuli, everyday interaction with others is much more dynamic, thus, conceivably harder to predict and decode, e.g., requiring integration across larger time windows (e.g. Arnal & Giraud, 2012; see Section 4.2). Diminished temporal integration abilities have indeed been reported in ASD for dynamic changes in auditory stimuli (reviewed in Kargas et al., 2015) and for sentence integration as reflected by deficient cross-cortical functional connectivity (Just et al., 2004). It is proposed here that similar deficits should be observed for decoding social dynamics. Figure 4, Panel B, depicts a metaphoric interpretation for how a system that lacks predictive coding ability at sensory level might fail to properly parse a dynamic social interaction into meaningful perceptual elements that make up an everyday sequence of social exchange. If predictions are lacking about “what” is likely to happen “when”, then processing of a dynamic sequence could either get stuck at a particular element or blend into an incomprehensible mix of sensory perceptions. Deficiencies in dynamic decoding should be prominently related to low frequency aberrations (delta, theta) and associated PAC, while more static stimuli could reveal deficient PAC (e.g. Khan et al., 2013) in relation to higher alpha/beta frequencies (Table 4, Hypothesis 7).

High-level social interactions require complex predictions that depend on a larger number of factors and conditions and tend to be context-specific rather than generally applicable across situations. For instance, a quite complex coordination of social signals (facial, postural), situational and conversational context factors, including memory of past events, may lead to the prediction that a particular remark might be ironic (“Wonderful weather here in Britain, isn’t it?!”) – or if the prediction is not in place, that the adjustment of the meaning (from actual to ironic) is successful based on a quite specific error-signal. In general the temporal integration windows required for optimal decoding of a social interaction are likely to increase with the complexity of the interaction, requiring ever finer coordination between low frequencies across cortical areas with local high frequency gamma (conforming to Arnal & Giraud, 2012). Recently, the relevance of theta-based networks for high-level social cognition (perspective taking, mentalizing) has indeed been reported (Bögels et al., 2014; Wang et al., 2016), and requires further investigation with ASD participants and finer-grained analysis of local phase and cross-frequency coupling. We hypothesise (Table 4, Hypothesis 8) that low frequency (delta-theta) cross-cortical phase coupling should be affected in ASD in conjunction with reduced local PAC and possibly inter-trial phase coherence during high-level social cognition.

Another aspect that ties in with this notion of deficient predictive coding of complex social dynamics is the consistent reports of ASD-specific aberrations of the amygdala, which has been tied to processing of emotional aspects of social interactions (Baron-Cohen et al., 2000; Kleinhans et al., 2016). This extends the notion of predictive coding failures due to complex visual dynamics in ASD to include complex emotional dynamics. If a system cannot adequately predict what others might feel or what the system itself might feel in response to an action or stimulus, then the interaction of the system with others will be hampered. In conclusion, social stimuli and contexts are arguably harder and more complex to predict than events that do not involve human factors, thus, future research might reveal that it is this specific characteristic of social situations that makes them harder to process in ASD rather than their inherent “quality” of being “social”. For example, this would be consistent with recent suggestions of the role of multiple bottom-up visual cues in “split second social perception” (Freeman & Johnson, 2016).

**Figure 4.**
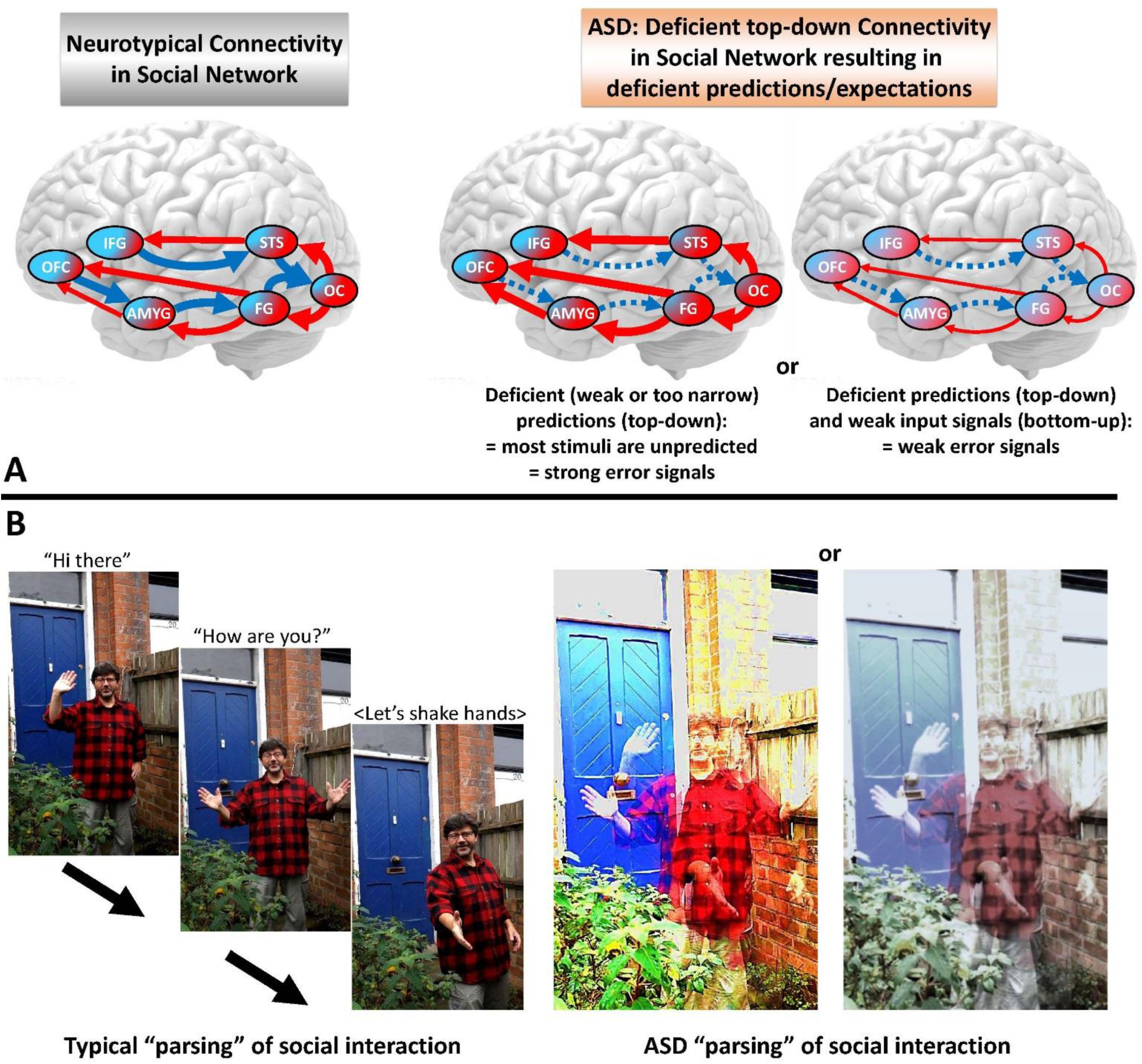
Proposed processing in the social network and hypothetical percepts. Conforming to Fig. 2, the left column shows assumed neurotypical connectivity and percepts, while the middle column shows the case hypothesised for hyper-sensitivities and the right column for hypo-sensitivities. Panel A depicts global connectivity in a hypothetical social network, where nodes of the network have been adapted from McPartland and Jeste (2015) and are employed as an illustration rather than a veridical model. In agreement with the proposed general framework of ASD (see Fig. 2) top-down connectivity in lower frequencies is assumed to be affected, reducing the predictive precision of the ASD system for social stimuli and especially for dynamic social sequences and interactions. High-level social interaction in particular has been associated with predictive coding in theta frequencies (Bögels et al., 2015), which is proposed to be affected in ASD. A metaphoric illustration of how a social sequence could be misprocessed in ASD is given in Panel B (middle and right). The left-hand side of Panel B shows how a neurotypical system might parse a quick sequence of movements and utterances into meaningful elements. We propose that in the ASD system predictive coding of “what” is expected to happen “when” is affected to an extent that elements cannot be effectively separated into meaningful chunks. Either elements are blended together (artistic impressions shown here in the middle and on the right) or processing could “get stuck” at a certain element that captures processing resources and impedes sequence processing to proceed (not explicitly depicted, but would imply that the sequence shown on the left is interrupted at an early element). Importantly, the more unusual (infrequent) a certain social dynamic would be, the harder the decoding would become – true for a neurotypical system, for an ASD system even more so. Note that the current example merely serves the purpose of visualising our hypotheses; we do not propose that all individuals with ASD will have difficulties decoding this particular greeting sequence (as it is very common and prototypical) or that their subjective experience matches the shown “blended” sequence.

## 7. Further refinements and future focus

The proposed framework implies that future research should continue to focus on long-range brain connectivity (structural and functional) involving the frontal cortex, with a particular emphasis on directionality and phase coupling in lower brain frequencies that should also be employed to measure cross-frequency coupling with gamma at the local level (e.g. Khan et al., 2013). The latter would provide an optimal reflection of regulation-efficiency of local processing, including efficiency of top-down regulatory influences and feedback for predictive coding at the local level. This is consistent with the proposal that low- and mid-range frequencies (delta-theta, alpha-beta) might provide top-down temporal integration windows that are optimal for predicting “when” things may happen, while higher beta frequencies may code for “what is likely to happen (Arnal & Giraud, 2012), and with gamma subcycles coding for “what” has actually happened (e.g. Mehta, Lee, & Wilson, 2002) as well as reflecting the current (mis-) match with a prediction in form of a feedforward error signal (Arnal & Giraud, 2012). Throughout Section 6 testable hypotheses have been derived based on the proposed framework that are summarised in Table 4 and may guide future investigation of oscillopathies based on those novel methods that have been discussed (Khan et al., 2013). The latter would provide an optimal reflection of regulation-efficiency of local processing, including efficiency of top-down regulatory influences and feedback for predictive coding at the local level. This is consistent with the proposal that low- and mid-range frequencies (delta-theta, alpha-beta) might provide top-down temporal integration windows that are optimal for predicting “when” things may happen, while higher beta frequencies may code for “what is likely to happen (Arnal & Giraud, 2012), and with gamma subcycles coding for “what” has actually happened (e.g. as well as reflecting the current (mis-) match with a prediction in form of a feedforward error signal.

Future research should therefore aim for refinement of the profiling of oscillatory cortical networks, perhaps using graph theory metrics (e.g. Sporns, 2003; Stam & Reijneveld, 2007) in addition to the metrics discussed above, with the aim of producing a connectivity ‘fingerprint’ which could characterise individual differences in terms of local/global connectivity patterns and also in degree of resting state vs. sensory responsivity. Connectivity research employing EEG/MEG should also extend beyond the cortex to establish whether oscillopathies can be observed in the function of subcortical structures (Simon & Wallace, 2016; Uddin, 2015), using localisation techniques suited for deep electromagnetic sources (Mills et al., 2012). The proposed predictive coding framework implies that resting state measures, reflecting the absence of specific predictions about the world, should be combined with measures of specific stimulation input in autism research, allowing analysis of predictive coding optimisation (or lack thereof) as reflected in long-range phase synchronisation and local cross-frequency coupling in relation to stimuli. This framework also allows comparing predictive coding deficits in relation to social and non-social stimuli, potentially shedding light on commonalities and/or on a continuum of differences.

There needs to be a greater focus in ASD cohorts on individual differences in symptom severity and symptom patterns, together with the inclusion of different groups more representative of different ages and levels of functioning. A clearer definition of the sensory/perceptual and the social phenotype should inform task and participant selection, with subsequent detailed statistical mapping of the relationship between the atypical behavioural characteristics and the wider oscillatory profile. In particular, ASD subjects reporting hypo-sensitivity to sensory stimuli requires further investigation with regards to the framework developed in this review (see Table 4, Hypothesis 3). Overall, this work could assist the development of more discriminatory ASD biomarkers, possibly identifying subtypes within the diagnostic category and/or endophenotypes and link to ongoing genetics research.

Additionally, translational work in this area offers the possibility of identifying potential pharmacological interventions with normalising of GBA, PAC, long-range connectivity and other atypical oscillatory signatures as a measure of effectiveness (Politte, Henry, & McDougle, 2014; Tyzio et al., 2014). Direct manipulation of neurophysiological activity, for example using repetitive TMS, has been shown to be effective in modulating various aspects of ASD symptomatology, including repetitive behaviour (Baruth et al., 2011), motor control and perceptual binding (Casanova et al., 2015). This aspect of research into brain oscillations, then, could take forward the possibility of developing interventions and thus not only contribute to a better understanding of the condition but also to its amelioration.

## Acknowledgments

Robert Seymour was sponsored by a Cotutelle PhD Scholarship awarded by Aston University and Macquarie University.

